# MetAMDB: Metabolic Atom Mapping Database

**DOI:** 10.1101/2021.10.05.463172

**Authors:** Collin Starke, Andre Wegner

## Abstract

MetAMDB (https://metamdb.tu-bs.de/) is an open source metabolic atom mapping database, providing atom mappings for around 75000 metabolic reactions. Each atom mapping can be inspected and downloaded either as a RXN file or as a graphic in SVG format. In addition, MetAMDB offers the possibility of automatically creating atom mapping models based on user-specified metabolic networks. These models can be of any size (small to genome scale) and can subsequently be used in standard ^13^C metabolic flux analysis software.

## Introduction

Metabolic fluxes are a valuable read-out for biomedical research [1] as well as for biotechnological applications [2]. However, metabolic fluxes are not measurable and must be inferred through metabolic modeling. Besides constraint-based modeling [3], the most successful approaches are based on stable isotope labeling experiments [4, 5]. Here, a stable isotope (mostly ^13^C) labeled substrate is applied and the metabolization of this compound will lead to specific isotopic enrichment patterns in downstream metabolites. For example, ^13^C metabolic flux analysis combines experimentally determined isotopic enrichment patterns of metabolites with a metabolic atom mapping model to infer metabolic fluxes (Antoniewicz, 2018). An atom mapping describes the one-to-one correspondence between a substrate atom and a product atom in a metabolic reaction [6]. Thus, within an atom mapping model one can follow every atom throughout the metabolic network. The creation of such a model is a time consuming process and requires detailed knowledge about the reaction mechanisms, and is therefore, mostly limited to smaller simplified models of metabolism. There are a few resources available that provide atom mappings for single reactions which are either restricted in access [7] or solely computationally derived [8]. There is currently no resource available that can automatically generate custum atom mapping models. This probably has to do with the fact that even if the atom mappings of the individual model reactions are known, it is difficult to combine them, because the ordering of atoms across all reactions must be consistent.

Here, we present MetAMDB (Metabolic Atom Mapping Database), a freely accessible, web-resource for metabolic atom mappings (https://metamdb.tu-bs.de/). Atom mappings for individual reactions can be inspected and downloaded either as a RXN file or as a graphic in SVG format. In addition, users can submit custom metabolic models and download the corresponding carbon atom mapping models with a single click. As such, MetAMDB will greatly facilitate the use of bigger (up to genome scale) metabolic atom mapping models, as they can be generated now easily, even by non-experts. The MetAMDB source code is available at https://github.com/CollinStark/metamdb under the MIT licence.

### Data Collection and Processing

The general overview of how we collected, processed, and curated the data included in MetAMDB is depicted in Figure 1.

**Figure 1:**
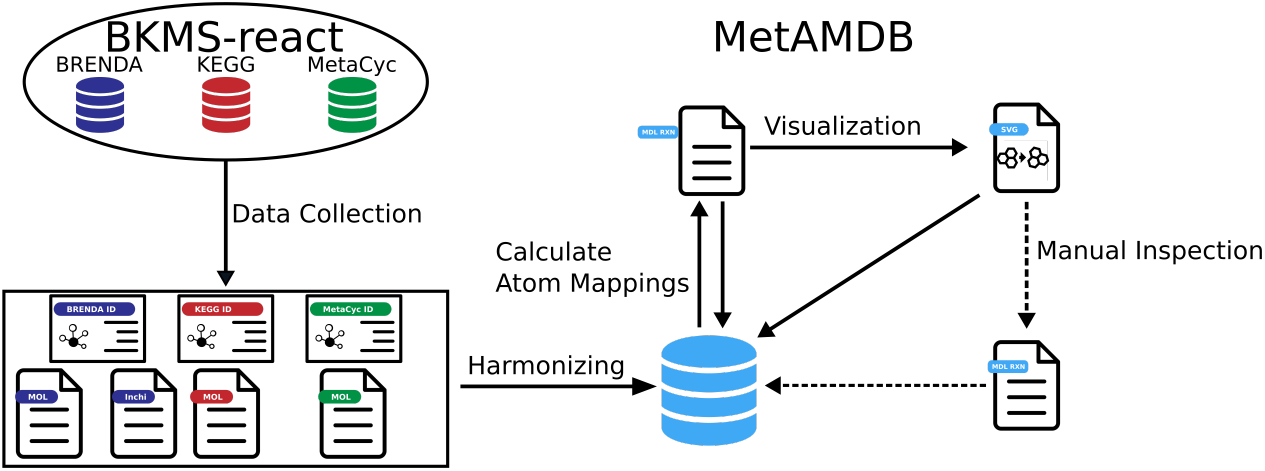
All data were collected from BKMS-react. To avoid duplications, metabolites were matched internally with a synonym algorithm and identical reactions were linked to a single reaction in MetAMDB. With all reaction data available, atom mappings for each reaction were calculated and visualized. Afterwards, a set of core reactions (around 1000) that cover central carbon metabolism were curated.

### Collection of Reaction and Metabolite Data

To include a comprehensive set of reactions, we parsed reaction data from BKMS-react [9], a biochemical reaction database containing data mainly collected from the three databases BRENDA [10], KEGG [11], and MetaCyc [7]. Since all of these databases are using different identifiers, we initially linked reaction data from each database to a single reaction in the MetAMDB database. Additionally, we collected metabolite data from BKMS-react and mapped each of the metabolites to their original database. To avoid duplications, we matched metabolites internally with a synonym algorithm and combined metabolites with different protonation states as this does not affect atom mappings. With this approach, we linked multiple reaction and metabolite identifiers of external databases to a single entry in the MetAMDB database. This harmonization is advantageous to simplify the creation of atom mapping models because identifiers from different databases can be used. We have decided to consider reactions in isolation from organisms, because we assume that an enzyme with the same substrates and products that is present in multiple organisms will have identical atom mappings. Finally, we parsed metabolite structures in the form of MOL files from BRENDA. If BRENDA MOL files were not available, we considered MetaCyc or KEGG, depending on their availability.

### Automated Generation of Atom Mappings

After we harmonized the individual reactions from BRENDA, KEGG, and MetaCyc to one unique reaction in MetAMDB, we calculated atom mappings for all of these reactions. Atom mappings track every substrate atom in a reaction to their corresponding product atom. The difficulty of generating an atom mapping, is therefore, dependent on the complexity of the corresponding reaction. Many different atom mapping algorithms have been developed, like DREAM [12], CLCA [13], Pathway Tools Software [14], Reaction Decoder Tool (RDT) [15], and a recently developed automatic mapper [16], with respective advantages and disadvantages for particular enzyme classes [6]. We decided to use RDT for the automatic generation of atom mappings because RDT reached the overall highest accuracy in the study by Gonzalez et. al. [6]. In addition, RDT is open-source, which makes it easy to modify and integrate in existing pipelines. RDT generates atom mappings in the form of RXN files from unmapped MOL files.

### Curation of Atom Mappings

RDT has a reported accuracy of around 90% (with some enzyme categories being better than others) [6], meaning approximately 10% of mappings contain an error. If one would look at a single reaction, this seems like an acceptable error rate, but it turns out to be rather problematic in a series of reactions such as a metabolic model. In a metabolic model, even one small error in a single reaction can lead to completely incorrect results. Figure 2A depicts a simplified reaction scheme of glycolysis and the pyruvate dehydrogenase complex. Suppose, we perform a stable isotope labeling experiment with [1-^13^C_1_]glucose as a tracer. With correct atom mappings, we can follow the labeled carbon atom to acetyl-CoA. However, if there is an atom mapping error in just one of the 10 glycolytic reactions, then the simulated labeling can drastically change. For example, an atom mapping error in the aldolase reaction (Figure 2B) would move the ^13^C labeled atom away from acetyl-CoA to the carbon dioxide released by the pyruvate dehydrogenase complex (Figure 2A).

**Figure 2:**
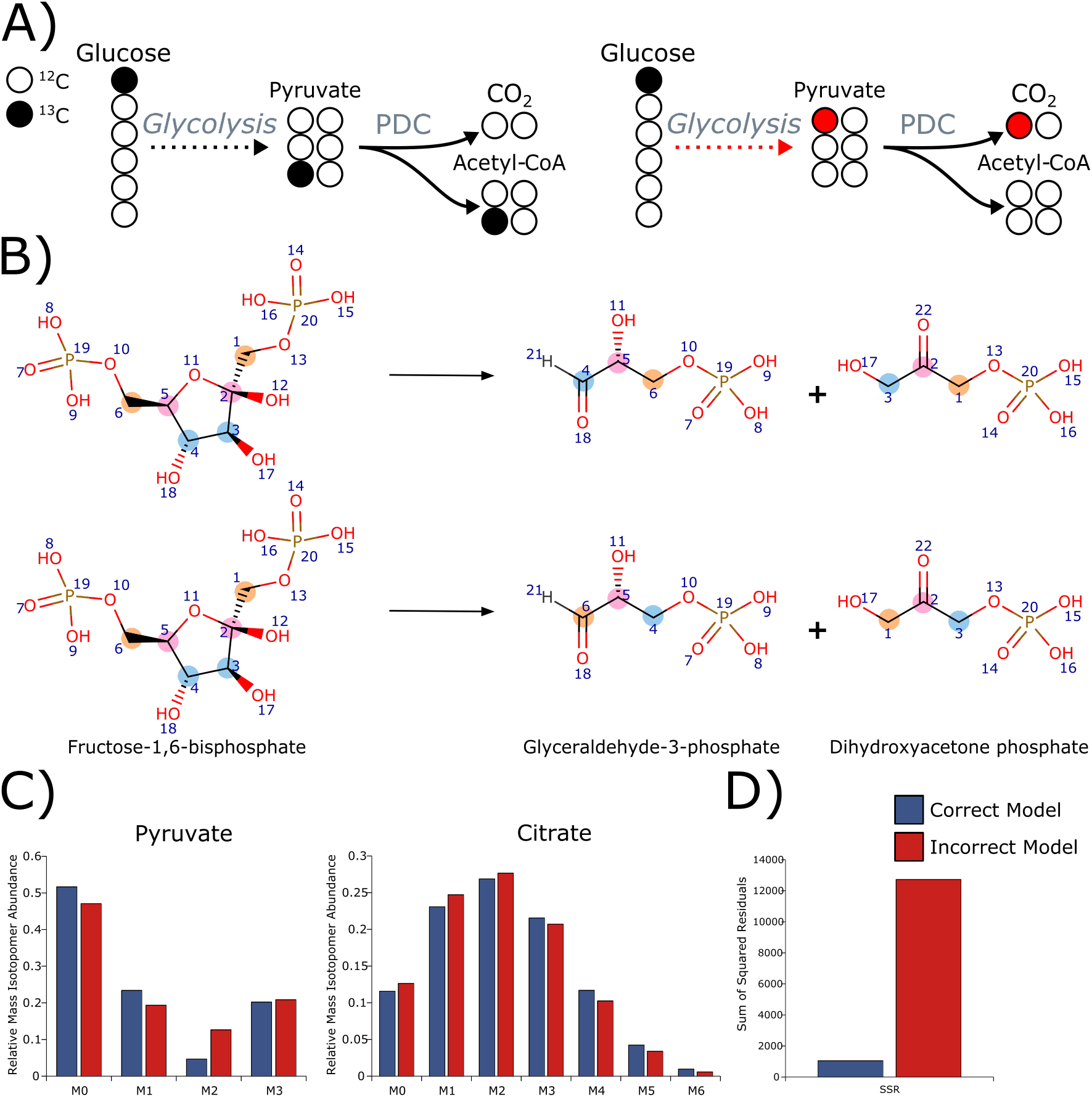
(**A**) Glycolysis converts glucose into two molecules of pyruvate, which in turn is converted to acetyl-CoA and CO_2_. [1-^13^C_1_]glucose will label the third pyruvate carbon ([3-^13^C_1_]pyruvate). The first carbon is split off in the pyruvate dehydrogenase complex resulting in a labeled acetyl-CoA ([2-^13^C_1_]acetyl-CoA). A single error in glycolysis will lead to inaccuracies in all downstream reactions. In this example, the aldolase atom mapping is incorrect, leading to [1-^13^C_1_] pyruvate instead of [3-^13^C_1_]pyruvate. This error then continues in the pyruvate dehydrogenase reaction, leading to labeling in carbon dioxide and not in acetyl-CoA. (**B**) In the aldolase reaction, fructose-1,6-bisphosphate is converted to glyceraldehyde-3-phosphate and dihydroxyacetone phosphate. The 1st and 6th carbon atom of fructose-1,6-bisphosphate are the phosphate carbons in their respective metabolites. In the second reaction an error is introduced that swaps the 1st and 3rd carbon atom of each metabolite, resulting in the 3rd and 4th carbon being the phosphate carbon. (**C**) Simulated MIDs for pyruvate and citrate for two different atom mapping models using the flux set with the best fit (Table S5). The correct model (blue) includes the correct aldolase atom mapping whereas the incorrect model (red) includes the wrong atom mapping of the aldolase reaction described above. (**D**) The sum of squared residuals (SSR) for the correct model was about 10x lower than the SSR of the incorrect model.

To demonstrate the impact of such an error on the estimation of intracellular metabolic fluxes, we performed ^13^C metabolic flux analysis using the INCA software suite [17]. To that end, we used the *E.coli* model of central carbon metabolism provided by INCA [18] and compared the simulated isotopic enrichment patterns, sum-of-squared residuals, and flux distribution of the original model to the model containing the atom mapping error in the aldolase reaction (see Supplementary File 1 for method details). We evaluated the isotopic enrichment in terms of mass isotopomer distribution (MID) vectors. MIDs describe the fractional abundance of each isotopologue normalized to the sum of all possible isotopologues [19]. Using a mixture of 75% [1-^13^C_1_] and 25% [U-^13^C_6_] labeled glucose as a tracer, the incorrect atom mapping model largely overestimates the M2 of pyruvate (Figure 2C) and subsequently the pyruvate dehydrogenase flux (Table S5). Overall, the estimated fluxes using the incorrect atom mapping model have much broader 95 % confidence intervalls (Table S5) and a much higher sum of squared residuals (Figure 2D), showing that the incorrect atom mapping model does not fit the data well. These results clearly demonstrate that error-free atom mapping models are fundamental for correct flux estimations. For this reason, we decided to curate a set of core reactions (around 1000) that cover central carbon metabolism. Of these, approximately 6 % contained an error. It is also possible for MetAMDB users to curate atom mappings, which will be integrated into MetAMDB after review.

### Automatic Generation of Atom Mapping Models

Atom mapping models are generated from user-submitted metabolic models and database atom mappings (Figure 3A). The metabolic model format is a custom csv-based format containing four columns: (1) reaction name, (2) substrates, (3) reversibility, (4) and products. Identifiers from either BRENDA, KEGG, or MetaCyc must be specified in square brackets for the reaction name, substrates, and products in order to match the reaction correctly to a MetAMDB reaction (Figure 3A). More details can be found in the MetAMDB documentation (https://collinstark.github.io/metamdb-docs/getting-started/). Database mappings are put into the ABC-Format after each metabolite. The ABC-Format is a per element atom mapping representation, which for now can only display carbon mappings in MetAMDB (e.g. (Glutamine (abcde) −> Glutamate (abcde)). While the ABC-Format lacks additional information compared to RXN files, like atom-specific characteristics and bond types, it makes up for it with its simplicity and human readability. For the example above, the first glutamine carbon (“a”) maps to the first glutamate carbon (“a”), and subsequently to the first carbon of alpha-ketoglutarate.

**Figure 3:**
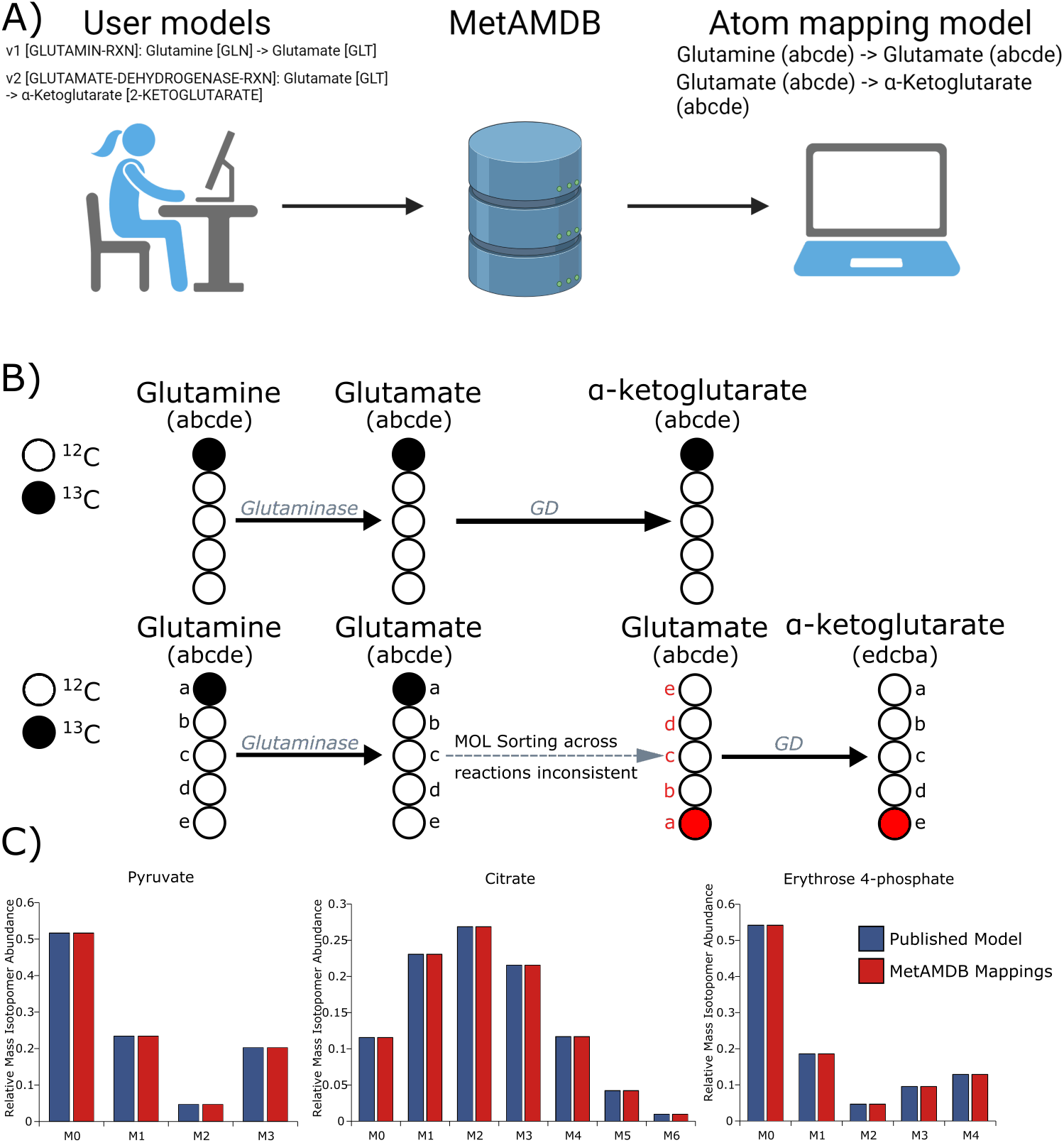
(**A**) Users can upload custom metabolic models to automatically generate an atom mapping model. The metabolic models must include a reaction identifier and metabolite identifiers in square brackets either from BRENDA, KEGG, or MetaCyc. In this example MetaCyc identifiers are used. The output is an atom mapping model in the abc format. (**B**) Glutamine is converted by glutaminase to glutamate and subsequently to alpha-ketoglutarate by glutamate dehydrogenase (GD). The carbon backbone in these reactions is unchanged. However, correct atom mappings can still result in errors in the atom mapping model. For example, if the ordering of the glutamate carbons of the second reaction (glutamate dehydrogenase) is transposed compared to the ordering in the first reaction (glutaminase), the first carbon of glutamine will be mapped to the fifth carbon of alpha-ketoglutarate, even though the atom mappings for the two single reactions are correct. (**C**) Simulation of isotopic enrichment for the metabolites pyruvate, citrate, and erythrose 4-phosphate using a published atom mapping model ([18], blue) and MetAMDB derived atom mappings for the same metabolic network (red).

One would assume that the automatic generation of this carbon atom mapping model is straightforward, as the carbon backbone in these reactions is unchanged (Figure 3B). However, to automatically generate a correct atom mapping model, the atoms of a compound have to be ordered identically across all MOL files. For example, if the ordering of the glutamate carbons in the glutamate dehydrogenase reaction is transposed compared to the ordering in the glutaminase reaction, the first carbon of glutamine would be mapped to the fifth carbon of alpha-ketoglutarate (Figure 3B). Theoretically, this problem should be fixed by ordering techniques like the IUPAC numbering. However, most mapping algorithms like RDT do not employ such strategies, making user-generated models by a beginner even more error-prone. To overcome this problem, we developed a sorting algorithm that orders each metabolite by its IUPAC numbering. Another problem for the automated generation of atom mapping models are symmetric metabolites (e.g. succinate), as symmetries will lead to multiple equivalent atom mappings. Currently, we only added symmetries for the curated reactions. A symmetric atom will be marked with a period symbol (“.”) in the RXN file, while the generated image indicates symmetries with duplicate atom numbers (Figure S1). A symmetric molecule in an atom mapping model will result in multiple mappings for the same metabolite.

To show that MetAMDB can automatically generate correct atom mapping models, we again used the *E.coli* model of central carbon metabolism provided by INCA [18] (see Table S1). First, we converted this model into a model readable by MetAMDB by adding MetaCyc identifiers (Table S4). To compare the atom mapping model from MetAMDB to the original model, we limited the number of carbons for acetyl-CoA to the acetyl moiety and removed symmetries as they were not present in the original model. However, the atom mapping models still cannot be compared directly because MetAMDB uses a different atom sorting. For that reason, we decided to compare the simulated isotopic enrichment patterns for both atom mapping models using INCA. As we have shown above, a small atom mapping error will lead to significant differences in the isotopic enrichment patterns (Figure 2C). Using the same flux set (Table S5) for both models, we obtained identical isotopic enrichment patterns for all metabolites included in the metabolic network (Figure 3C, Table S6, and Table S8), showing that MetAMDB can automatically generate correct atom mapping models.

### Database Content and Usage

MetAMDB provides atom mappings for reactions present in BKMS-react [9]. Every atom mapping is linked to their respective compounds and enzymes, and all of them are accessible by their KEGG, MetaCyc, or BRENDA identifiers. MetAMDB atom mappings are freely available on the public webpage (https://metamdb.tu-bs.de/). The database consists of approximately 75000 reactions with atom mappings generated by RDT and corresponding structural images in SVG format. Around 1000 of these reactions were curated. Each atom mapping, if possible, is linked to a BRENDA, KEGG, and MetaCyc reaction. The database contains about 94000 metabolites with MOL files and structural images. To find a specific atom mapping, users can search MetAMDB using different database queries. For example, reaction identifiers (BRENDA, KEGG, MetaCyc) or reaction names can be used. Users can also search for specific subtrates and products included in a reaction. We have implemented a fuzzy string search, meaning that glucose as a search term will also match reactions that include glucose-6-phosphate. A successful query will show the reaction name, the identifier, substrates and products and whether the atom mapping was curated. Additional information is contained on the reaction page itself, which can be opened from the database query results. The atom mapping can be viewed (or downloaded) in the RXN file format. An image of the reaction structure, which includes the atom mapping can also be opened and downloaded. In addition to the atom mappings, compounds of the reaction can also be inspected. The compound page has additional information such as InChIs, InChIKeys, and also MOL files that were used in the generation of the atom mapping. A help page with documentation for possible use cases can be found on the MetAMDB website.

Users can upload custom metabolic models to obtain an atom mapping model for their specified reactions. Metabolic models require database identifiers for the reaction as well as the substrates and products. If necessary, reactions can be simpliefied by omitting substrates or products from the reaction ( for example, cofactors such as ADP and ATP). A successfully parsed metabolic model can be viewed on the reaction model page, which displays reactions and compounds with the subsequent linked MetAMDB pages. Atom mappings in the abc-format are also displayed, which if necessary can be manually adjusted. If a manual or simplified reaction is needed (e.g. multiple linear reactions condensed into one), reactions with custom identifiers can be written that require manual atom mappings. The final model can be downloaded and used for further analysis and modelling applications.

Some of the functionality of MetAMDB can also be accessed through a REST-API. For example reaction and pathway data are accessible, as well as the database search.

## Conclusion

The quantitative analysis of metabolism is an important part of systems biology. These analysis are often based on stable isotope labeled experiments and require an atom-resolved understanding of the metabolic network. That means one has to be able to follow the fate of every atom in the network. Creating such a model is a time-cosuming and error-prone process, in particular for larger models. To overcome this problem, we developed MetAMDB. MetAMDB provides atom mappings for single reactions but also supports the automatic generation of atom mapping models. To our knowledge, this is the first implementation of such a feature. As such, MetAMDB will greatly facilitate the use of bigger (up to genome scale) metabolic atom mapping models, as they can be generated now easily, even by non-experts. Since a single error in the atom mappings can make the entire downstream analysis faulty, we have curated a subset of reactions of central carbon metabolism and and have integrated the possibility in MetAMDB that users can correct incorrect atom mappings.

## Acknowledgements

The authors acknowledge funding by the German Research Foundation (DFG, Project-ID 34509606 – TRR 51), the MWK of Lower Saxony (SMART BIOTECS alliance between the Technische Universität Braunschweig and the Leibniz Universität Hannover) and the German Federal Ministry of Education and Research (BMBF, PeriNAA - 01ZX1916B).

## Notes

### Competing Interest Statement

The authors have declared no competing interest.

