## Supplement for "MetAMDB: Metabolic Atom Mapping Database"

This document contains the following information in support of the primary article:

### Method Text: Flux estimation and MID simulation

All flux estimations and MID simulations were done in INCA (Young, 2014) for the provided *E. coli* model (Young et al., 2008). The model is a dynamic, nonsteady-state model and was used with its predetermined settings by INCA. For the flux set, fluxes were estimated and for the confidence interval calculation the INCA parameter continuation was used. Since the model is dynamic, multiple MIDs were simulated but only the last time point was used in this paper (0.0178).

*Table S1:* ***Atom mapping model of the INCA E. coli model (Young, 2014).***

| ID | Substrates | | Products |
|---|---|---|---|
| R1 | G6P (abcdef) | <-> | F6P (abcdef) |
| R2 | F6P (abcdef) | -> | FBP (abcdef) |
| R3 | FBP (abcdef) | <-> | DHAP (cba) + GAP (def) |
| R4 | DHAP (abc) | <-> | GAP (abc) |
| R5 | GAP (abc) | <-> | PG3 (abc) |
| R6 | PG3 (abc) | <-> | PEP (abc) |
| R7 | PEP (abc) | -> | Pyr (abc) |
| R8 | G6P (abcdef) | -> | PG6 (abcdef) |
| R9 | PG6 (abcdef) | -> | Ru5P (bcdef) + CO2 (a) |
| R10 | Ru5P (abcde) | <-> | X5P (abcde) |
| R11 | Ru5P (abcde) | <-> | R5P (abcde) |
| R12 | X5P (abcde) | <-> | GAP (cde) + EC2 (ab) |
| R13 | F6P (abcdef) | <-> | E4P (cdef) + EC2 (ab) |
| R14 | S7P (abcdefg) | <-> | R5P (cdefg) + EC2 (ab) |
| R15 | F6P (abcdef) | <-> | GAP (def) + EC3 (abc) |
| R16 | S7P (abcdefg) | <-> | E4P (defg) + EC3 (abc) |
| R17 | PG6 (abcdef) | -> | KDPG (abcdef) |
| R18 | KDPG (abcdef) | -> | Pyr (abc) + GAP (def) |
| R19 | Pyr (abc) | -> | AcCoA (bc) + CO2 (a) |
| R20 | OAA (abcd) + AcCoA (ef) | -> | Cit (dcbfea) |
| R21 | Cit (abcdef) | <-> | ICit (abcdef) |
| R22 | ICit (abcdef) | <-> | AKG (abcde) + CO2 (f) |
| R23 | AKG (abcde) | -> | SucCoA (bcde) + CO2 (a) |
| R24 | SucCoA (abcd) | <-> | Suc (abcd) |
| R25 | Suc (abcd) | <-> | Fum (abcd) |
| R26 | Fum (abcd) | <-> | Mal (abcd) |
| R27 | Mal (abcd) | <-> | OAA (abcd) |
| R28 | Mal (abcd) | -> | Pyr (abc) + CO2 (d) |
| R29 | PEP (abc) + CO2 (d) | <-> | OAA (abcd) |
| R30 | AcCoA (ab) | <-> | Ac (ab) |
| R31 | DHAP (abc) | <-> | Glyc3P (abc) |
| R32 | Glyc3P (abc) | -> | Glyc (abc) |
| R33 | Glyc (abc) | -> | HPA (abc) |
| R34 | HPA (abc) | -> | PDO (abc) |
| R35 | AKG (abcde) | -> | Glu (abcde) |
| R36 | Glu (abcde) | -> | Gln (abcde) |
| R37 | Glu (abcde) | -> | Pro (abcde) |
| R38 | Glu (abcde) + CO2 (f) + Gln (ghijk) + Asp (lmno) + AcCoA (pq) | -> | Arg (abcdef) + AKG (ghijk) + Fum (lmno) + Ac (pq) |
| R39 | OAA (abcd) + Glu (efghi) | -> | Asp (abcd) + AKG (efghi) |

| | | | |
|---|---|---|---|
| R40 | Asp (abcd) | -> | Asn (abcd) |
| R41 | Pyr (abc) + Glu (defgh) | -> | Ala (abc) + AKG (defgh) |
| R42 | PG3 (abc) + Glu (defgh) | -> | Ser (abc) + AKG (defgh) |
| R43 | Ser (abc) | <-> | Gly (ab) + MEETHF (c) |
| R44 | Gly (ab) | <-> | CO2 (a) + MEETHF (b) |
| R45 | Thr (abcd) | -> | Gly (ab) + AcCoA (cd) |
| R46 | Ser (abc) + AcCoA (de) | -> | Cys (abc) + Ac (de) |
| R47 | Asp (abcd) + Pyr (efg) + Glu (hijkl) + SucCoA (mnop) | -> | LL_DAP (abcdgfe) + AKG (hijkl) + Suc (mnop) |
| R48 | LL_DAP (abcdefg) | -> | Lys (abcdef) + CO2 (g) |
| R49 | Asp (abcd) | -> | Thr (abcd) |
| R50 | Asp (abcd) + METHF (e) + Cys (fgh) + SucCoA (ijkl) | -> | Met (abcde) + Pyr (fgh) + Suc (ijkl) |
| R51 | Pyr (abc) + Pyr (def) + Glu (ghijk) | -> | Val (abcef) + CO2 (d) + AKG (ghijk) |
| R52 | AcCoA (ab) + Pyr (cde) + Pyr (fgh) + Glu (ijklm) | -> | Leu (abdghe) + CO2 (c) + CO2 (f) + AKG (ijklm) |
| R53 | Thr (abcd) + Pyr (efg) + Glu (hijkl) | -> | Ile (abfcdg) + CO2 (e) + AKG (hijkl) |
| R54 | PEP (abc) + PEP (def) + E4P (ghij) + Glu (klmno) | -> | Phe (abcefghij) + CO2 (d) + AKG (klmno) |
| R55 | PEP (abc) + PEP (def) + E4P (ghij) + Glu (klmno) | -> | Tyr (abcefghij) + CO2 (d) + AKG (klmno) |
| R56 | Ser (abc) + R5P (defgh) + PEP (ijk) + E4P (lmno) + PEP (pqr) + Gln (stuvw) | -> | Trp (abcedklmnoj) + CO2 (i) + GAP (fgh) + Pyr (pqr) + Glu (stuvw) |
| R57 | R5P (abcde) + FTHF (f) + Gln (ghijk) + Asp (lmno) | -> | His (edcbaf) + AKG (ghijk) + Fum (lmno) |
| R58 | MEETHF (a) | -> | METHF (a) |
| R59 | MEETHF (a) | -> | FTHF (a) |
| R60 | Gluc.pre (abcdef) | -> | G6P (abcdef) |
| R61 | Gluc.ext (abcdef) | -> | G6P (abcdef) |
| R62 | Cit.ext (abcdef) | -> | Cit (abcdef) |
| R63 | Glyc.ext (abc) + Dummy.ext | <-> | Glyc (abc) + Dummy |
| R64 | PDO (abc) | -> | PDO.ext (abc) |
| R65 | Ac (ab) | -> | Ac.ext (ab) |
| R66 | CO2 (a) | -> | CO2.ext (a) |
| R67 | 0.488*Ala + 0.281*Arg + 0.229*Asn + 0.229*Asp + 0.087*Cys + 0.25*Glu + 0.25*Gln + 0.582*Gly + 0.09*His + 0.276*Ile + 0.428*Leu + 0.326*Lys + 0.146*Met + 0.176*Phe + 0.21*Pro + 0.205*Ser + 0.241*Thr + 0.054*Trp + 0.131*Tyr + 0.402*Val + 0.205*G6P + 0.071*F6P + 0.754*R5P + 0.129*GAP + | -> | 39.68*Biomass |

| | | | |
|---|---|---|---|
| | 0.619*PG3 + 0.051*PEP + 0.083*Pyr + 2.51*AcCoA + 0.087*AKG + 0.34*OAA + 0.443*MEETHF | | |
| R68 | Dummy | -> | Dummy.ext |

*Table S2:* ***Atom mapping model of the INCA E. coli model (Young, 2014) with an error in R3.***

In this model we introduced an error in the aldolase reaction with the ID R3. Both dihydroxyacetone phosphate's (DHAP) and glyceraldehyde 3-phosphate's (GAP) mappings are mirrored, meaning the first and third carbon atoms are swapped in each metabolite.

| ID | Substrates | | Products |
|---|---|---|---|
| R1 | G6P (abcdef) | <-> | F6P (abcdef) |
| R2 | F6P (abcdef) | -> | FBP (abcdef) |
| **R3** | **FBP (abcdef)** | **<->** | **DHAP (abc) + GAP (fed)** |
| R4 | DHAP (abc) | <-> | GAP (abc) |
| R5 | GAP (abc) | <-> | PG3 (abc) |
| R6 | PG3 (abc) | <-> | PEP (abc) |
| R7 | PEP (abc) | -> | Pyr (abc) |
| R8 | G6P (abcdef) | -> | PG6 (abcdef) |
| R9 | PG6 (abcdef) | -> | Ru5P (bcdef) + CO2 (a) |
| R10 | Ru5P (abcde) | <-> | X5P (abcde) |
| R11 | Ru5P (abcde) | <-> | R5P (abcde) |
| R12 | X5P (abcde) | <-> | GAP (cde) + EC2 (ab) |
| R13 | F6P (abcdef) | <-> | E4P (cdef) + EC2 (ab) |
| R14 | S7P (abcdefg) | <-> | R5P (cdefg) + EC2 (ab) |
| R15 | F6P (abcdef) | <-> | GAP (def) + EC3 (abc) |
| R16 | S7P (abcdefg) | <-> | E4P (defg) + EC3 (abc) |
| R17 | PG6 (abcdef) | -> | KDPG (abcdef) |
| R18 | KDPG (abcdef) | -> | Pyr (abc) + GAP (def) |
| R19 | Pyr (abc) | -> | AcCoA (bc) + CO2 (a) |
| R20 | OAA (abcd) + AcCoA (ef) | -> | Cit (dcbfea) |
| R21 | Cit (abcdef) | <-> | ICit (abcdef) |
| R22 | ICit (abcdef) | <-> | AKG (abcde) + CO2 (f) |
| R23 | AKG (abcde) | -> | SucCoA (bcde) + CO2 (a) |
| R24 | SucCoA (abcd) | <-> | Suc (abcd) |
| R25 | Suc (abcd) | <-> | Fum (abcd) |
| R26 | Fum (abcd) | <-> | Mal (abcd) |
| R27 | Mal (abcd) | <-> | OAA (abcd) |
| R28 | Mal (abcd) | -> | Pyr (abc) + CO2 (d) |
| R29 | PEP (abc) + CO2 (d) | <-> | OAA (abcd) |
| R30 | AcCoA (ab) | <-> | Ac (ab) |
| R31 | DHAP (abc) | <-> | Glyc3P (abc) |
| R32 | Glyc3P (abc) | -> | Glyc (abc) |
| R33 | Glyc (abc) | -> | HPA (abc) |
| R34 | HPA (abc) | -> | PDO (abc) |
| R35 | AKG (abcde) | -> | Glu (abcde) |
| R36 | Glu (abcde) | -> | Gln (abcde) |
| R37 | Glu (abcde) | -> | Pro (abcde) |
| R38 | Glu (abcde) + CO2 (f) + Gln (ghijk) + Asp (lmno) + AcCoA (pq) | -> | Arg (abcdef) + AKG (ghijk) + Fum (lmno) + Ac (pq) |

| | | | |
|---|---|---|---|
| R39 | OAA (abcd) + Glu (efghi) | -> | Asp (abcd) + AKG (efghi) |
| R40 | Asp (abcd) | -> | Asn (abcd) |
| R41 | Pyr (abc) + Glu (defgh) | -> | Ala (abc) + AKG (defgh) |
| R42 | PG3 (abc) + Glu (defgh) | -> | Ser (abc) + AKG (defgh) |
| R43 | Ser (abc) | <-> | Gly (ab) + MEETHF (c) |
| R44 | Gly (ab) | <-> | CO2 (a) + MEETHF (b) |
| R45 | Thr (abcd) | -> | Gly (ab) + AcCoA (cd) |
| R46 | Ser (abc) + AcCoA (de) | -> | Cys (abc) + Ac (de) |
| R47 | Asp (abcd) + Pyr (efg) + Glu (hijkl) + SucCoA (mnop) | -> | LL_DAP (abcdgfe) + AKG (hijkl) + Suc (mnop) |
| R48 | LL_DAP (abcdefg) | -> | Lys (abcdef) + CO2 (g) |
| R49 | Asp (abcd) | -> | Thr (abcd) |
| R50 | Asp (abcd) + METHF (e) + Cys (fgh) + SucCoA (ijkl) | -> | Met (abcde) + Pyr (fgh) + Suc (ijkl) |
| R51 | Pyr (abc) + Pyr (def) + Glu (ghijk) | -> | Val (abcef) + CO2 (d) + AKG (ghijk) |
| R52 | AcCoA (ab) + Pyr (cde) + Pyr (fgh) + Glu (ijklm) | -> | Leu (abdghe) + CO2 (c) + CO2 (f) + AKG (ijklm) |
| R53 | Thr (abcd) + Pyr (efg) + Glu (hijkl) | -> | Ile (abfcdg) + CO2 (e) + AKG (hijkl) |
| R54 | PEP (abc) + PEP (def) + E4P (ghij) + Glu (klmno) | -> | Phe (abcefghij) + CO2 (d) + AKG (klmno) |
| R55 | PEP (abc) + PEP (def) + E4P (ghij) + Glu (klmno) | -> | Tyr (abcefghij) + CO2 (d) + AKG (klmno) |
| R56 | Ser (abc) + R5P (defgh) + PEP (ijk) + E4P (lmno) + PEP (pqr) + Gln (stuvw) | -> | Trp (abcedklmnoj) + CO2 (i) + GAP (fgh) + Pyr (pqr) + Glu (stuvw) |
| R57 | R5P (abcde) + FTHF (f) + Gln (ghijk) + Asp (lmno) | -> | His (edcbaf) + AKG (ghijk) + Fum (lmno) |
| R58 | MEETHF (a) | -> | METHF (a) |
| R59 | MEETHF (a) | -> | FTHF (a) |
| R60 | Gluc.pre (abcdef) | -> | G6P (abcdef) |
| R61 | Gluc.ext (abcdef) | -> | G6P (abcdef) |
| R62 | Cit.ext (abcdef) | -> | Cit (abcdef) |
| R63 | Glyc.ext (abc) + Dummy.ext | <-> | Glyc (abc) + Dummy |
| R64 | PDO (abc) | -> | PDO.ext (abc) |
| R65 | Ac (ab) | -> | Ac.ext (ab) |
| R66 | CO2 (a) | -> | CO2.ext (a) |
| R67 | 0.488*Ala + 0.281*Arg + 0.229*Asn + 0.229*Asp + 0.087*Cys + 0.25*Glu + 0.25*Gln + 0.582*Gly + 0.09*His + 0.276*Ile + 0.428*Leu + 0.326*Lys + 0.146*Met + 0.176*Phe + 0.21*Pro + 0.205*Ser + 0.241*Thr + 0.054*Trp + 0.131*Tyr + 0.402*Val + 0.205*G6P + 0.071*F6P + | -> | 39.68*Biomass |

| | | | |
|---|---|---|---|
| | 0.754*R5P + 0.129*GAP + 0.619*PG3 + 0.051*PEP + 0.083*Pyr + 2.51*AcCoA + 0.087*AKG + 0.34*OAA + 0.443*MEETHF | | |
| R68 | Dummy | -> | Dummy.ext |

*Table S3:* ***Atom mapping model of E. coli (see Table S1) with MetAMDB atom mappings.***

We utilized the E. coli atom mapping model in combination with MetAMDB reaction ids to generate MetAMDB atom mappings. Most model reactions are simplified, which means most atom mappings will stay similar to the original ones. Some atom mappings were manually adjusted, which mainly include reduction of Acetyl-CoA and Succinyl-CoA. Simplified atom mappings only specify two and four carbon atoms respectively, while database atom mappings include all carbon atom in the mapping.

| ID | Substrates | | Products |
|---|---|---|---|
| R1 | G6P (abcdef) | <-> | F6P (abcdef) |
| R2 | F6P (abcdef) | -> | FBP (abcdef) |
| R3 | FBP (abcdef) | <-> | DHAP (abc) + GAP (def) |
| R4 | DHAP (abc) | <-> | GAP (cba) |
| R5 | GAP (abc) | <-> | PG3 (abc) |
| R6 | PG3 (abc) | <-> | PEP (abc) |
| R7 | PEP (abc) | -> | Pyr (abc) |
| R8 | G6P (abcdef) | -> | PG6 (abcdef) |
| R9 | PG6 (abcdef) | -> | Ru5P (bcdef) + CO2 (a) |
| R10 | Ru5P (abcde) | <-> | X5P (abcde) |
| R11 | Ru5P (abcde) | <-> | R5P (abcde) |
| R12 | X5P (abcde) | <-> | GAP (cde) + EC2 (ab) |
| R13 | F6P (abcdef) | <-> | E4P (cdef) + EC2 (ab) |
| R14 | S7P (abcdefg) | <-> | R5P (cdefg) + EC2 (ab) |
| R15 | F6P (abcdef) | <-> | GAP (def) + EC3 (abc) |
| R16 | S7P (abcdefg) | <-> | E4P (defg) + EC3 (abc) |
| R17 | PG6 (abcdef) | -> | KDPG (abcdef) |
| R18 | KDPG (abcdef) | -> | Pyr (abc) + GAP (def) |
| R19 | Pyr (abc) | -> | AcCoA (bc) + CO2 (a) |
| R20 | OAA (abcd) + AcCoA (tu) | -> | Cit (tubcda) |
| R21 | Cit (abcdef) | <-> | ICit (abcdef) |
| R22 | ICit (abcdef) | <-> | AKG (edcba) + CO2 (f) |
| R23 | AKG (abcde) | -> | SucCoA (edcb) + CO2 (a) |
| R24 | SucCoA (fghi) | <-> | Suc (fghi) |
| R25 | Suc (abcd) | <-> | Fum (abcd) |
| R26 | Fum (abcd) | <-> | Mal (abcd) |
| R27 | Mal (abcd) | <-> | OAA (abcd) |
| R28 | Mal (abcd) | -> | Pyr (abc) + CO2 (d) |
| R29 | PEP (abc) + CO2 (d) | <-> | OAA (abcd) |
| R30 | AcCoA (ab) | <-> | Ac (ab) |
| R31 | DHAP (abc) | <-> | Glyc3P (abc) |
| R32 | Glyc3P (abc) | -> | Glyc (cba) |
| R33 | Glyc (abc) | -> | HPA (abc) |
| R34 | HPA (abc) | -> | PDO (abc) |
| R35 | AKG (abcde) | -> | Glu (abcde) |
| R36 | Glu (abcde) | -> | Gln (abcde) |

| | | | |
|---|---|---|---|
| R37 | Glu (abcde) | -> | Pro (abcde) |
| R38 | Glu (abcde) + CO2 (f) + Gln (ghijk) + Asp (lmno) + AcCoA (pq) | -> | Arg (abcdef) + AKG (ghijk) + Fum (lmno) + Ac (pq) |
| R39 | OAA (abcd) + Glu (efghi) | -> | Asp (abcd) + AKG (efghi) |
| R40 | Asp (abcd) | -> | Asn (abcd) |
| R41 | Pyr (abc) + Glu (defgh) | -> | Ala (abc) + AKG (defgh) |
| R42 | PG3 (abc) + Glu (defgh) | -> | Ser (abc) + AKG (defgh) |
| R43 | Ser (abc) | <-> | Gly (ab) + MEETHF (c) |
| R44 | Gly (ab) | <-> | CO2 (a) + MEETHF (b) |
| R45 | Thr (abcd) | -> | Gly (ab) + AcCoA (cd) |
| R46 | Ser (abc) + AcCoA (de) | -> | Cys (abc) + Ac (de) |
| R47 | Asp (abcd) + Pyr (efg) + Glu (hijkl) + SucCoA (mnop) | -> | LL_DAP (abcdgfe) + AKG (hijkl) + Suc (mnop) |
| R48 | LL_DAP (abcdefg) | -> | Lys (abcdef) + CO2 (g) |
| R49 | Asp (abcd) | -> | Thr (abcd) |
| R50 | Asp (abcd) + METHF (e) + Cys (fgh) + SucCoA (ijkl) | -> | Met (abcde) + Pyr (fgh) + Suc (ijkl) |
| R51 | Pyr (abc) + Pyr (def) + Glu (ghijk) | -> | Val (abcef) + CO2 (d) + AKG (ghijk) |
| R52 | AcCoA (ab) + Pyr (cde) + Pyr (fgh) + Glu (ijklm) | -> | Leu (abdghe) + CO2 (c) + CO2 (f) + AKG (ijklm) |
| R53 | Thr (abcd) + Pyr (efg) + Glu (hijkl) | -> | Ile (abfcdg) + CO2 (e) + AKG (hijkl) |
| R54 | PEP (abc) + PEP (def) + E4P (ghij) + Glu (klmno) | -> | Phe (abcefghij) + CO2 (d) + AKG (klmno) |
| R55 | PEP (abc) + PEP (def) + E4P (ghij) + Glu (klmno) | -> | Tyr (abcefghij) + CO2 (d) + AKG (klmno) |
| R56 | Ser (abc) + R5P (defgh) + PEP (ijk) + E4P (lmno) + PEP (pqr) + Gln (stuvw) | -> | Trp (abcedklmnoj) + CO2 (i) + GAP (fgh) + Pyr (pqr) + Glu (stuvw) |
| R57 | R5P (abcde) + FTHF (f) + Gln (ghijk) + Asp (lmno) | -> | His (edcbaf) + AKG (ghijk) + Fum (lmno) |
| R58 | MEETHF (a) | -> | METHF (a) |
| R59 | MEETHF (a) | -> | FTHF (a) |
| R60 | Gluc.pre (abcdef) | -> | G6P (abcdef) |
| R61 | Gluc.ext (abcdef) | -> | G6P (abcdef) |
| R62 | Cit.ext (abcdef) | -> | Cit (abcdef) |
| R63 | Glyc.ext (abc) + Dummy.ext | <-> | Glyc (abc) + Dummy |
| R64 | PDO (abc) | -> | PDO.ext (abc) |
| R65 | Ac (ab) | -> | Ac.ext (ab) |
| R66 | CO2 (a) | -> | CO2.ext (a) |
| R67 | 0.488*Ala + 0.281*Arg + 0.229*Asn + 0.229*Asp + 0.087*Cys + 0.25*Glu + 0.25*Gln + 0.582*Gly + 0.09*His + 0.276*Ile + 0.428*Leu + 0.326*Lys + 0.146*Met + | -> | 39.68*Biomass |

| | | | |
|---|---|---|---|
| | 0.176*Phe + 0.21*Pro + 0.205*Ser + 0.241*Thr + 0.054*Trp + 0.131*Tyr + 0.402*Val + 0.205*G6P + 0.071*F6P + 0.754*R5P + 0.129*GAP + 0.619*PG3 + 0.051*PEP + 0.083*Pyr + 2.51*AcCoA + 0.087*AKG + 0.34*OAA + 0.443*MEETHF | | |
| R68 | Dummy | -> | Dummy.ext |

*Table S4:* ***Metabolic model of E. coli (see Table S1) with MetaCyc identifiers***

| reaction | substrates | arrow | products |
|---|---|---|---|
| v1 [PGLUCISOM-RXN] | G6P [Glucose-6-phosphate] | <-> | F6P [CPD-18719] |
| v2 [6PFRUCTPHOS-RXN] | F6P [CPD-18719] | -> | FBP [FRUCTOSE-16-DIPHOSPHATE] |
| v3 [F16ALDOLASE-RXN] | FBP [FRUCTOSE-16-DIPHOSPHATE] | <-> | DHAP [136411] + GAP [GAP] |
| v4 [TRIOSEPISOMERIZATION-RXN] | DHAP [136411] | <-> | GAP [GAP] |
| v5 | GAP (abc) [GAP] | <-> | PG3 (abc) |
| v6 | PG3 (abc) | <-> | PEP (abc) [PHOSPHO-ENOL-PYRUVATE] |
| v7 [PEPDEPHOS-RXN] | PEP [PHOSPHO-ENOL-PYRUVATE] | -> | Pyr [PYRUVATE] |
| v8 | G6P (abcdef) | -> | PG6 (abcdef) [CPD-2961] |
| v9 [RXN-9952] | PG6 [CPD-2961] | -> | Ru5P [RIBULOSE-5P] + CO2 [CARBON-DIOXIDE] |
| v10 [RIBULP3EPIM-RXN] | Ru5P [RIBULOSE-5P] | <-> | X5P [XYLULOSE-5-PHOSPHATE] |
| v11 [RIB5PISOM-RXN] | Ru5P [RIBULOSE-5P] | <-> | R5P [C00117] |
| v12 | X5P (abcde) [XYLULOSE-5-PHOSPHATE] | ↔ | EC2 (ab) + GAP (cde) [GAP] |
| v13 | F6P (abcdef) [CPD-18719] | ↔ | EC2 (ab) + E4P (cdef) |
| v14 | S7P (abcdefg) [D-SEDOHEPTULOSE-7-P] | ↔ | EC2 (ab) + R5P (cdefg) [C00117] |
| v15 | F6P (abcdef) [CPD-18719] | ↔ | EC3 (abc) + GAP (def) [GAP] |
| v16 | S7P (abcdefg) [D-SEDOHEPTULOSE-7-P] | ↔ | EC3 (abc) + E4P (defg) |
| v17 | PG6 [CPD-2961] (abcdef) | -> | KDPG [2-DEHYDRO-3-DEOXY-D-GLUCONATE] (abcdef) |
| v18 [KDPGALDOL-RXN] | KDPG [2-KETO-3-DEOXY-6-P-GLUCONATE] | -> | Pyr [PYRUVATE] + GAP [GAP] |
| v19 [PYRUVDEH-RXN] | Pyr [PYRUVATE] | -> | AcCoA [ACETYL-COA] + CO2 [CARBON-DIOXIDE] |
| v20 [CITSYN-RXN] | OAA [OXALACETIC_ACID] + AcCoA [ACETYL-COA] | -> | Cit [CIT] |
| v21 | Cit [CIT] (abcdef) | <-> | Icit [C00311] (abcdef) |
| v22 [ISOCITRATE-DEHYDROGENASE-NAD+-RXN] | Icit [C00311] | <-> | AKG [2-KETOGLUTARATE] + CO2 [CARBON-DIOXIDE] |
| v23 [2OXOGLUTARATEDEH- | AKG [2-KETOGLUTARATE] | -> | SucCoA [SUC-COA] + CO2 [CARBON-DIOXIDE] |

| | | | |
|---|---|---|---|
| RXN] | | | |
| v24 [SUCCCOASYN-RXN] | SucCoA [SUC-COA] | <-> | Suc [SUC] |
| v25 [SUCC-FUM-OXRED-RXN] | Suc [SUC] | <-> | Fum [FUM] |
| v26 [FUMHYDR-RXN] | Fum [FUM] | <-> | Mal [MAL] |
| v27 [MALATE-DEH-RXN] | Mal [MAL] | <-> | OAA [OXALACETIC_ACID] |
| v28 [MALIC-NADP-RXN] | Mal [MAL] | -> | Pyr [PYRUVATE] + CO2 [CARBON-DIOXIDE] |
| v29 [PEPCARBOX-RXN] | PEP [PHOSPHO-ENOL-PYRUVATE] + CO2 [HCO3] | <-> | OAA [OXALACETIC_ACID] |
| v30 [ACETYL-COA-HYDROLASE-RXN] | AcCoA [ACETYL-COA] | <-> | Ac [ACET] |
| v31 [GLYC3PDEHYDROGBIOSYN-RXN] | DHAP [136411] | <-> | Glyc3P [C00093] |
| v32 [RXN-14965] | Glyc3P [GLYCEROL-3P] | -> | Glyc [GLYCEROL] |
| v33 | Glyc (abc) | -> | HPA (abc) |
| v34 | HPA (abc) | -> | PDO (abc) |
| v35 | AKG (abcde) [2-KETOGLUTARATE] | -> | Glu (abcde) |
| v36 | Glu (abcde) | -> | Gln (abcde) |
| v37 | Glu (abcde) | -> | Pro (abcde) |
| v38 | Glu (abcde) + CO2 (f) + Gln (ghijk) + Asp (lmno) + AcCoA (pq) | -> | Arg (abcdef) + AKG (ghijk) [2-KETOGLUTARATE] + Fum (lmno) + Ac (pq) |
| v39 | OAA (abcd) + Glu (efghi) | -> | Asp (abcd) + AKG (efghi) [2-KETOGLUTARATE] |
| v40 | Asp (abcd) | -> | Asn (abcd) |
| v41 | Pyr (abc) + Glu (defgh) | -> | Ala (abc) + AKG (defgh) [2-KETOGLUTARATE] |
| v42 | PG3 (abc) + Glu (defgh) | -> | Ser (abc) + AKG (defgh) [2-KETOGLUTARATE] |
| v43 | Ser (abc) | <-> | Gly (ab) + MEETHF (c) |
| v44 | Gly (ab) | <-> | CO2 (a) + MEETHF (b) |
| v45 | Thr (abcd) | -> | Gly (ab) + AcCoA (cd) |
| v46 | Ser (abc) + AcCoA (de) | -> | Cys (abc) + Ac (de) |
| v47 | Asp (abcd) + Pyr (efg) + Glu (hijkl) + SucCoA (mnop) | -> | LL_DAP (abcdgfe) + AKG (hijkl) + Suc (mnop) |
| v48 | LL_DAP (abcdefg) | -> | Lys (abcdef) + CO2 (g) |
| v49 | Asp (abcd) | -> | Thr (abcd) |
| v50 | Asp (abcd) + METHF (e) + Cys (fgh) + SucCoA (ijkl) | -> | Met (abcde) + Pyr (fgh) + Suc (ijkl) |
| v51 | Pyr (abc) + Pyr (def) + Glu (ghijk) | -> | Val (abcef) + CO2 (d) + AKG (ghijk) |
| v52 | AcCoA (ab) + Pyr (cde) + Pyr (fgh) + Glu (ijklm) | -> | Leu (abdghe) + CO2 (c) + CO2 (f) + AKG (ijklm) |
| v53 | Thr (abcd) + Pyr (efg) + Glu | -> | Ile (abfcdg) + CO2 (e) + AKG |

| | | | |
|---|---|---|---|
| | (hijkl) | | (hijkl) |
| v54 | PEP (abc) + PEP (def) + E4P (ghij) + Glu (klmno) | -> | Phe (abcefghij) + $CO_2$ (d) + AKG (klmno) |
| v55 | PEP (abc) + PEP (def) + E4P (ghij) + Glu (klmno) | -> | Tyr (abcefghij) + $CO_2$ (d) + AKG (klmno) |
| v56 | Ser (abc) + R5P (defgh) + PEP (ijk) + E4P (lmno) + PEP (pqr) + Gln (stuvw) | -> | Trp (abcedklmnoj) + $CO_2$ (i) + GAP (fgh) + Pyr (pqr) + Glu (stuvw) |
| v57 | R5P (abcde) + FTHF (f) + Gln (ghijk) + Asp (lmno) | -> | His (edcbaf) + AKG (ghijk) + Fum (lmno) |
| v58 | MEETHF (a) | -> | METHF (a) |
| v59 | MEETHF (a) | -> | FTHF (a) |
| v60 | Gluc.pre (abcdef) | -> | G6P (abcdef) |
| v61 | Gluc.ext (abcdef) | -> | G6P (abcdef) |
| v62 | Cit.ext (abcdef) | -> | Cit (abcdef) |
| v63 | Glyc.ext (abc) + Dummy.ext | <-> | Glyc (abc) + Dummy |
| v64 | PDO (abc) | -> | PDO.ext (abc) |
| v65 | Ac (ab) | -> | Ac.ext (ab) |
| v66 | $CO_2$ (a) [CARBON-DIOXIDE] | -> | $CO_2$.ext (a) |
| v67 | 0.488*Ala + 0.281*Arg + 0.229*Asn + 0.229*Asp + 0.087*Cys + 0.25*Glu + 0.25*Gln + 0.582*Gly + 0.09*His + 0.276*Ile + 0.428*Leu + 0.326*Lys + 0.146*Met + 0.176*Phe + 0.21*Pro + 0.205*Ser + 0.241*Thr + 0.054*Trp + 0.131*Tyr + 0.402*Val + 0.205*G6P + 0.071*F6P + 0.754*R5P + 0.129*GAP + 0.619*PG3 + 0.051*PEP + 0.083*Pyr + 2.51*AcCoA + 0.087*AKG + 0.34*OAA + 0.443*MEETHF | -> | 39.68*Biomass |
| v68 | Dummy | -> | Dummy.ext |

*Table S5:* ***Flux estimation results for the E. coli model (see Table S1 and S2).***

| | | | Cor. Model | | Incor. Model | |
|---|---|---|---|---|---|---|
| Reaction | | | Flux | 95% Conf. | Flux | 95% Conf. |
| G6P ↔ F6P | R1 | net | 63,9 | [ 52,4 , 72,3 ] | 77,1 | [ NaN , 86.2144 ] |
| | | exch | 0,0 | [ 0,0 , 99,5 ] | 291,1 | [ 242.1618 , 338.2041 ] |
| F6P → FBP | R2 | net | 86,2 | [ 79,2 , 92,9 ] | 76,4 | [ NaN , 86.1196 ] |
| FBP ↔ DHAP + GAP | R3 | net | 86,2 | [ 79,2 , 92,9 ] | 76,4 | [ NaN , 86.1196 ] |
| | | exch | 595,6 | [ 0,0 , Inf ] | 2,71E+02 | [ 0 , 4.2154e+06 ] |
| DHAP ↔ GAP | R4 | net | -41,1 | [ -48,5 , -33,9 ] | -42,0 | [ -52.0429 , -31.6098 ] |
| | | exch | 246,4 | [ 189,8 , 349,0 ] | 31,9 | [ 28.1662 , 35.8754 ] |
| GAP ↔ PG3 | R5 | net | 56,3 | [ 49,9 , 64,4 ] | 57,7 | [ 52.0042 , 64.5625 ] |
| | | exch | 802500,0 | [ 165,6 , Inf ] | 4.1395e+05 | [ 0 , Inf ] |
| PG3 ↔ PEP | R6 | net | 54,7 | [ 49,1 , 60,2 ] | 56,4 | [ 50.5624 , 63.1878 ] |
| | | exch | 5571,9 | [ 153,8 , Inf ] | 5.2318e+05 | [ 0 , Inf ] |
| PEP → Pyr | R7 | | 48,6 | [ 43,2 , 56,6 ] | 53,5 | [ 48.0505 , 60.1951 ] |
| G6P → PG6 | R8 | | 34,7 | [ 28,9 , 45,0 ] | 23,7 | [ 21.0883 , 26.5164 ] |
| PG6 → Ru5P + CO2 | R9 | | 34,6 | [ 28,6 , 45,5 ] | 2.6482e-06 | [ 0 , 1.5161 ] |
| Ru5P ↔ X5P | R10 | net | 22,4 | [ 18,5 , 30,7 ] | -0,6 | [ -0.6396 , 0.5453 ] |
| | | exch | 108,9 | [ 0,0 , Inf ] | 4.7306e+06 | [ 9.3512e+04 , NaN ] |
| Ru5P ↔ R5P | R11 | net | 12,2 | [ 10,3 , 16,5 ] | 0,6 | [ NaN , 1.0447 ] |
| | | exch | 0,0 | [ 0,0 , Inf ] | 4.7306e+06 | [ 6.0445e+04 , NaN ] |
| X5P ↔ EC2 + GAP | R12 | net | 22,4 | [ 18,5 , 30,7 ] | -0,6 | [ -0.6396 , 0.5453 ] |
| | | exch | 3125,7 | [ 47,2 , Inf ] | 4.7306e+06 | [ 6.1687e+04 , NaN ] |
| F6P ↔ EC2 + E4P | R13 | net | -11,0 | [ -15,2 , -9,1 ] | 0,4 | [ -0.2091 , 0.4801 ] |
| | | exch | 5,8 | [ 0,0 , 13,3 ] | 61,3 | [ 49.7521 , 76.1032 ] |
| S7P ↔ EC2 + R5P | R14 | net | -11,4 | [ -15,7 , -9,4 ] | 0,1 | [ -0.3514 , 0.1613 ] |
| | | exch | 127,3 | [ 0,0 , Inf ] | 2.9505e+06 | [ 1.7499e+04 , Inf ] |
| F6P ↔ EC3 + GAP | R15 | net | -11,4 | [ -15,7 , -9,4 ] | 0,1 | [ -0.3514 , 0.1613 ] |

| | | | | | | | | | | | | | | |
|---|---|---|---|---|---|---|---|---|---|---|---|---|---|---|
| | | exch | 0,0 | [ | 0,0 | ’ | Inf | ] | 2.2132<br>e+06 | [ | 7.3161<br>e+03 | , | Inf | ] |
| S7P ↔ EC3 + E4P | R16 | net | 11,4 | [ | 9,4 | ’ | 15,7 | ] | -0,1 | [ | -<br>0.1613 | ’ | 0.3514 | ] |
| | | exch | 0,0 | [ | 0,0 | ’ | Inf | ] | 2.9505<br>e+06 | [ | 1.7138<br>e+04 | , | 2.9505<br>e+06 | ] |
| PG6 → KDPG | R17 | | 0,1 | [ | 0,0 | ’ | 0,9 | ] | 23,7 | [ | NaN | ’ | 26.714<br>8 | ] |
| KDPG → Pyr + GAP | R18 | | 0,1 | [ | 0,0 | ’ | 0,9 | ] | 23,7 | [ | NaN | ’ | 26.714<br>8 | ] |
| Pyr → AcCoA + CO2 | R19 | | 49,5 | [ | 44,2 | ’ | 57,7 | ] | 75,2 | [ | NaN | ’ | 84.433<br>0 | ] |
| OAA + AcCoA → Cit | R20 | | 46,6 | [ | 41,1 | ’ | 51,8 | ] | 72,7 | [ | 65.441<br>3 | , | 81.609<br>8 | ] |
| Cit ↔ Icit | R21 | net | 46,8 | [ | 41,5 | ’ | 54,5 | ] | 72,9 | [ | 65.322<br>6 | ’ | NaN | ] |
| | | exch | 94,6 | [ | 45,8 | ’ | 238,2 | ] | 1.2877<br>e+04 | [ | 172.23<br>10 | , | NaN | ] |
| Icit ↔ AKG + CO2 | R22 | net | 46,8 | [ | 41,5 | ’ | 54,5 | ] | 72,9 | [ | 65.322<br>6 | ’ | NaN | ] |
| | | exch | 18474,<br>0 | [ | 47,9 | ’ | Inf | ] | 9.1376<br>e+03 | [ | 184.37<br>10 | , | Inf | ] |
| AKG → SucCoA + CO2 | R23 | | 45,9 | [ | 40,5 | ’ | 53,9 | ] | 72,1 | [ | 65.347<br>6 | , | 81.243<br>4 | ] |
| SucCoA ↔ Suc | R24 | net | 45,5 | [ | 40,0 | ’ | 53,4 | ] | 71,7 | [ | 64.768<br>3 | , | 80.533<br>8 | ] |
| | | exch | 451,7 | [ | 0,0 | , | Inf | ] | 85,7 | [ | 0 | , | Inf | ] |
| Suc ↔ Fum | R25 | net | 45,9 | [ | 40,5 | ’ | 53,9 | ] | 72,1 | [ | 65.347<br>6 | , | 81.243<br>4 | ] |
| | | exch | 147,0 | [ | 53,1 | ’ | 7453,4 | ] | 4.3404<br>e+05 | [ | 6.8337 | ’ | 4.3405<br>e+05 | ] |
| Fum ↔ Mal | R26 | net | 46,2 | [ | 40,8 | ’ | 54,1 | ] | 72,3 | [ | 64.639<br>7 | , | 81.629<br>1 | ] |
| | | exch | 50699<br>0,0 | [ | 196,3 | ’ | Inf | ] | 8.6711<br>e+05 | [ | 610.90<br>94 | , | Inf | ] |
| Mal ↔ OAA | R27 | net | 43,1 | [ | 37,7 | ’ | 48,3 | ] | 72,3 | [ | 64.677<br>2 | , | 81.565<br>1 | ] |
| | | exch | 198,2 | [ | 134,6 | ’ | 28107,<br>0 | ] | 9.3089<br>e+05 | [ | 477.61<br>49 | , | Inf | ] |
| Mal → Pyr + CO2 | R28 | | 3,2 | [ | 1,9 | ’ | 4,0 | ] | 2.6563<br>e-06 | [ | 0 | ’ | 0.3336 | ] |
| PEP + CO2 ↔ OAA | R29 | net | 5,4 | [ | 4,1 | , | 6,4 | ] | 2,2 | [ | 1.7204 | , | 2.5668 | ] |
| | | exch | 7,1 | [ | 5,3 | ’ | 9,1 | ] | 1.0000<br>e-07 | [ | 0 | ’ | 0.7548 | ] |
| AcCOA ↔ Ac | R30 | net | -0,1 | [ | -0,2 | ’ | -0,1 | ] | -0,1 | [ | -<br>0.1607 | ’ | 0.0071 | ] |
| | | exch | 0,0 | [ | 0,0 | ’ | Inf | ] | 3.6533<br>e+05 | [ | 0 | , | 3.6533<br>e+05 | ] |
| DHAP ↔ Glyc3P | R31 | net | 127,3 | [ | 114,0 | ’ | 140,2 | ] | 118,5 | [ | 100.24<br>78 | , | 137.59<br>95 | ] |
| | | exch | 30024<br>0,0 | [ | 0,0 | ’ | Inf | ] | 4,3 | [ | 0 | ’ | Inf | ] |
| Glyc3P → Glyc | R32 | | 127,3 | [ | 114,0 | ’ | 140,2 | ] | 118,5 | [ | 100.24<br>78 | , | 137.59<br>95 | ] |
| Glyc → HPA | R33 | | 129,3 | [ | 116,6 | , | 142,0 | ] | 120,5 | [ | NaN | , | 139.48 | ] |

| | | | | | | | | | | | | | | |
|---|---|---|---|---|---|---|---|---|---|---|---|---|---|---|
| HPA → PDO | R34 | | 129,3 | [ | 116,6 | , | 142,0 | ] | 120,5 | [ | NaN | , | 39 139.48 39 | ] |
| AKG → Glu | R35 | | 5,8 | [ | 5,0 | , | 6,5 | ] | 5,2 | [ | NaN | , | 5.9591 | ] |
| Glu → Gln | R36 | | 0,6 | [ | 0,5 | , | 0,7 | ] | 0,5 | [ | 0.4270 | , | 0.6033 | ] |
| Glu → Pro | R37 | | 0,2 | [ | 0,2 | , | 0,2 | ] | 0,2 | [ | 0.1329 | , | 0.1877 | ] |
| Glu + CO2 + Gln + Asp + AcCoA → Arg + AKG + Fum + Ac | R38 | | 0,3 | [ | 0,2 | , | 0,3 | ] | 0,2 | [ | 0.1778 | , | 0.2511 | ] |
| OAA + Glu → Asp + AKG | R39 | | 1,6 | [ | 1,4 | , | 1,8 | ] | 1,6 | [ | NaN | , | 1.7603 | ] |
| Asp → Asn | R40 | | 0,2 | [ | 0,2 | , | 0,2 | ] | 0,2 | [ | 0.1449 | , | 0.2047 | ] |
| Pyr + Glu → Ala + AKG | R41 | | 0,4 | [ | 0,4 | , | 0,5 | ] | 0,4 | [ | 0.3087 | , | 0.4362 | ] |
| PG3 + Glu → Ser + AKG | R42 | | 1,0 | [ | 0,8 | , | 1,1 | ] | 0,8 | [ | 0.6325 | , | 0.9426 | ] |
| Ser ↔ Gly + MEETHF | R43 | net | 0,6 | [ | 0,5 | , | 0,6 | ] | 0,4 | [ | NaN | , | 0.4946 | ] |
| | | exch | 0,8 | [ | 0,7 | , | 1,0 | ] | 1,7 | [ | 1.2927 | , | 1.9677 | ] |
| Gly ↔ CO2 + MEETHF | R44 | net | 0,0 | [ | 0,0 | , | 0,1 | ] | 0,1 | [ | 0.0659 | , | 0.1461 | ] |
| | | exch | 0,2 | [ | 0,1 | , | 0,2 | ] | 0,0 | [ | 0 | , | 0.0667 | ] |
| Thr → Gly + AcCoA | R45 | | 0,0 | [ | 0,0 | , | 0,1 | ] | 0,2 | [ | NaN | , | 0.2085 | ] |
| Ser + AcCoA → Cys + Ac | R46 | | 0,2 | [ | 0,2 | , | 0,2 | ] | 0,2 | [ | 0.1474 | , | 0.2082 | ] |
| Asp + Pyr + Glu + SucCoA → LL_DAP + AKG + Suc | R47 | | 0,3 | [ | 0,2 | , | 0,3 | ] | 0,3 | [ | 0.2062 | , | 0.2914 | ] |
| LL_DAP → Lys + CO2 | R48 | | 0,3 | [ | 0,2 | , | 0,3 | ] | 0,3 | [ | 0.2062 | , | 0.2914 | ] |
| Asp → Thr | R49 | | 0,5 | [ | 0,4 | , | 0,5 | ] | 0,6 | [ | 0.4212 | , | 0.6604 | ] |
| Asp + METHF + Cys + SucCoA → Met + Pyr + Suc | R50 | | 0,1 | [ | 0,1 | , | 0,1 | ] | 0,1 | [ | 0.0924 | , | 0.1305 | ] |
| Pyr + Pyr + Glu → Val + CO2 + AKG | R51 | | 0,4 | [ | 0,3 | , | 0,4 | ] | 0,3 | [ | 0.2543 | , | 0.3593 | ] |
| AcCoA + Pyr + Pyr + Glu → Leu + CO2 + CO2 + AKG | R52 | | 0,4 | [ | 0,3 | , | 0,4 | ] | 0,3 | [ | 0.2708 | , | 0.3825 | ] |
| Thr + Pyr + Glu → Ile + CO2 + AKG | R53 | | 0,2 | [ | 0,2 | , | 0,3 | ] | 0,2 | [ | 0.1746 | , | 0.2467 | ] |
| PEP + PEP + E4P + Glu → Phe + CO2 + AKG | R54 | | 0,2 | [ | 0,1 | , | 0,2 | ] | 0,1 | [ | 0.1113 | , | 0.1573 | ] |
| PEP + PEP + E4P + Glu → Tyr + CO2 + AKG | R55 | | 0,1 | [ | 0,1 | , | 0,1 | ] | 0,1 | [ | 0.0829 | , | 0.1171 | ] |
| Ser + R5P + PEP + E4P + PEP + Gln → Trp + CO2 + GAP + Pyr + Glu | R56 | | 0,0 | [ | 0,0 | , | 0,1 | ] | 0,0 | [ | 0.0342 | , | 0.0483 | ] |
| R5P + FTHF + Gln + Asp → His + AKG + Fum | R57 | | 0,1 | [ | 0,1 | , | 0,1 | ] | 0,1 | [ | 0.0569 | , | 0.0804 | ] |
| MEETHF → METHF | R58 | | 0,1 | [ | 0,1 | , | 0,1 | ] | 0,1 | [ | 0.0924 | , | 0.1305 | ] |
| MEETHF → FTHF | R59 | | 0,1 | [ | 0,1 | , | 0,1 | ] | 0,1 | [ | 0.0569 | , | 0.0804 | ] |
| Gluc.pre → G6P | R60 | | 6,4 | [ | 5,4 | , | 7,4 | ] | 2.6341 e-06 | [ | 0 | , | 0.0361 | ] |
| Gluc.ext → G6P | R61 | | 92,3 | [ | 86,0 | , | 98,5 | ] | 101,0 | [ | 91.298 7 | , | 110.98 25 | ] |

| | | | | | | | | | | | | | |
|---|---|---|---|---|---|---|---|---|---|---|---|---|---|
| Cit.ext → Cit | R62 | 0,3 | [ | 0,2 | , | 0,3 | ] | 0,2 | [ | NaN | , | 0.2270 | ] |
| Glyc.ext ↔ Glyc | R63 | 2,0 | [ | 1,8 | , | 2,2 | ] | 1.9852 | [ | 1.6451 | , | 2.2693 | ] |
| | | 45,8 | [ | 0,0 | , | Inf | ] | 1.5725 e+03 | [ | 0 | , | Inf | ] |
| PDO → PDO.ext | R64 | 129,3 | [ | 116,6 | , | 142,0 | ] | 120,5 | [ | NaN | , | 139.48 39 | ] |
| Ac → Ac.ext | R65 | 0,3 | [ | 0,3 | , | 0,4 | ] | 0,3 | [ | 0.2586 | , | 0.3563 | ] |
| CO2 → CO2.ext | R66 | 176,3 | [ | 158,3 | , | 205,1 | ] | 219,6 | [ | 197.84 61 | , | NaN | ] |
| Biomass formation | R67 | 0,9 | [ | 0,8 | , | 1,0 | ] | 0,8 | [ | 0.6326 | , | 0.8938 | ] |
| Dummy → Dummy.ext | R68 | 2,0 | [ | 1,8 | , | 2,2 | ] | 2,0 | [ | 1.6451 | , | 2.2693 | ] |

*Table S6:* ***Mass Isotopomer Distribution (MID) results for the E. coli model without errors (see Table S1).***

| Mass Isotopomer | M0 | M1 | M2 | M3 | M4 | M5 | M6 | M7 | M8 | M9 |
|---|---|---|---|---|---|---|---|---|---|---|
| AKG5 | 0.1781 | 0.2673 | 0.2766 | 0.1839 | 0.0728 | 0.0213 | | | | |
| AcCoA | 0.5497 | 0.2209 | 0.2294 | | | | | | | |
| Ala2 | 0.5617 | 0.2153 | 0.223 | | | | | | | |
| Ala3 | 0.5295 | 0.2285 | 0.0452 | 0.1967 | | | | | | |
| Asp2a | 0.4995 | 0.2993 | 0.2012 | | | | | | | |
| Asp2b | 0.4995 | 0.2993 | 0.2012 | | | | | | | |
| Asp3 | 0.3156 | 0.3617 | 0.2292 | 0.0935 | | | | | | |
| Asp4 | 0.2447 | 0.3257 | 0.2443 | 0.1349 | 0.0504 | | | | | |
| Cit6 | 0.1157 | 0.2308 | 0.2688 | 0.2156 | 0.117 | 0.0424 | 0.0097 | | | |
| E4P | 0.5423 | 0.1861 | 0.0469 | 0.0956 | 0.1291 | | | | | |
| Glu4 | 0.2276 | 0.3181 | 0.2802 | 0.1341 | 0.0401 | | | | | |
| Glu5 | 0.1794 | 0.267 | 0.2762 | 0.1834 | 0.0726 | 0.0213 | | | | |
| Gly1 | 0.726 | 0.274 | | | | | | | | |
| Gly2 | 0.6531 | 0.1417 | 0.2052 | | | | | | | |
| Ile5a | 0.1737 | 0.2686 | 0.2782 | 0.1849 | 0.0732 | 0.0214 | | | | |
| Ile5b | 0.1737 | 0.2686 | 0.2782 | 0.1849 | 0.0732 | 0.0214 | | | | |
| Leu5 | 0.1787 | 0.2669 | 0.2749 | 0.1847 | 0.0728 | 0.0221 | | | | |
| Mal4 | 0.2412 | 0.3244 | 0.2511 | 0.1349 | 0.0485 | | | | | |
| Met4a | 0.1916 | 0.3436 | 0.2812 | 0.1468 | 0.0368 | | | | | |
| Met4b | 0.1916 | 0.3436 | 0.2812 | 0.1468 | 0.0368 | | | | | |
| Met5 | 0.1486 | 0.2939 | 0.2763 | 0.1779 | 0.0836 | 0.0198 | | | | |
| Phe2 | 0.7178 | 0.068 | 0.2142 | | | | | | | |
| Phe8a | 0.1696 | 0.1841 | 0.2227 | 0.1498 | 0.1243 | 0.0737 | 0.051 | 0.0177 | 0.007 | |
| Phe8b | 0.1696 | 0.1841 | 0.2227 | 0.1498 | 0.1243 | 0.0737 | 0.051 | 0.0177 | 0.007 | |
| Phe9 | 0.1612 | 0.1832 | 0.1608 | 0.1706 | 0.1307 | 0.093 | 0.0529 | 0.03 | 0.0113 | 0.0063 |
| Pyr3 | 0.5169 | 0.2342 | 0.0467 | 0.2023 | | | | | | |
| Ser2a | 0.5055 | 0.3183 | 0.1762 | | | | | | | |
| Ser2b | 0.6885 | 0.1014 | 0.2101 | | | | | | | |
| Ser2c | 0.5055 | 0.3183 | 0.1762 | | | | | | | |
| Suc4 | 0.2376 | 0.323 | 0.2581 | 0.1348 | 0.0465 | | | | | |
| Thr3 | 0.3157 | 0.3616 | 0.2291 | 0.0935 | | | | | | |
| Thr4 | 0.2448 | 0.3257 | 0.2442 | 0.1349 | 0.0504 | | | | | |
| Tyr2 | 0.7179 | 0.068 | 0.2141 | | | | | | | |
| Val4 | 0.3024 | 0.2428 | 0.3009 | 0.1013 | 0.0526 | | | | | |
| Val5 | 0.2844 | 0.2429 | 0.1959 | 0.1752 | 0.0554 | 0.0464 | | | | |

*Table S7:* ***Mass Isotopomer Distribution (MID) results for the E. coli model with an aldolase error (see Table S2).***

| Mass Isotopomer | M0 | M1 | M2 | M3 | M4 | M5 | M6 | M7 | M8 | M9 |
|---|---|---|---|---|---|---|---|---|---|---|
| AKG5 | 0.1957 | 0.2761 | 0.2773 | 0.1687 | 0.0661 | 0.0162 | | | | |
| AcCoA | 0.5826 | 0.1785 | 0.2388 | | | | | | | |
| Ala2 | 0.6084 | 0.1681 | 0.2236 | | | | | | | |
| Ala3 | 0.5027 | 0.1837 | 0.1182 | 0.1953 | | | | | | |
| Asp2a | 0.5064 | 0.3101 | 0.1836 | | | | | | | |
| Asp2b | 0.5064 | 0.3101 | 0.1836 | | | | | | | |
| Asp3 | 0.3327 | 0.3728 | 0.2264 | 0.068 | | | | | | |
| Asp4 | 0.254 | 0.333 | 0.25 | 0.1305 | 0.0324 | | | | | |
| Cit6 | 0.1264 | 0.2473 | 0.2767 | 0.2072 | 0.1026 | 0.034 | 0.0058 | | | |
| E4P | 0.3974 | 0.2889 | 0.0608 | 0.1644 | 0.0884 | | | | | |
| Glu4 | 0.2496 | 0.3347 | 0.2557 | 0.1292 | 0.0308 | | | | | |
| Glu5 | 0.1963 | 0.2759 | 0.2771 | 0.1685 | 0.066 | 0.0162 | | | | |
| Gly1 | 0.71 | 0.29 | | | | | | | | |
| Gly2 | 0.636 | 0.1319 | 0.2321 | | | | | | | |
| Ile5a | 0.1939 | 0.2766 | 0.2779 | 0.1691 | 0.0662 | 0.0163 | | | | |
| Ile5b | 0.1939 | 0.2766 | 0.2779 | 0.1691 | 0.0662 | 0.0163 | | | | |
| Leu5 | 0.2194 | 0.2546 | 0.273 | 0.1655 | 0.067 | 0.0204 | | | | |
| Mal4 | 0.254 | 0.3331 | 0.25 | 0.1305 | 0.0325 | | | | | |
| Met4a | 0.2331 | 0.3608 | 0.2703 | 0.1154 | 0.0204 | | | | | |
| Met4b | 0.2331 | 0.3608 | 0.2703 | 0.1154 | 0.0204 | | | | | |
| Met5 | 0.178 | 0.3094 | 0.2748 | 0.1663 | 0.0618 | 0.0097 | | | | |
| Phe2 | 0.6693 | 0.0853 | 0.2454 | | | | | | | |
| Phe8a | 0.178 | 0.1748 | 0.1937 | 0.1942 | 0.1148 | 0.0853 | 0.0402 | 0.0136 | 0.0053 | |
| Phe8b | 0.178 | 0.1748 | 0.1937 | 0.1942 | 0.1148 | 0.0853 | 0.0402 | 0.0136 | 0.0053 | |
| Phe9 | 0.1706 | 0.165 | 0.1352 | 0.2052 | 0.1316 | 0.085 | 0.0654 | 0.0246 | 0.0123 | 0.005 |
| Pyr3 | 0.4709 | 0.1937 | 0.1267 | 0.2087 | | | | | | |
| Ser2a | 0.5536 | 0.3078 | 0.1386 | | | | | | | |
| Ser2b | 0.6469 | 0.1166 | 0.2365 | | | | | | | |
| Ser2c | 0.5536 | 0.3078 | 0.1386 | | | | | | | |
| Suc4 | 0.254 | 0.3331 | 0.25 | 0.1305 | 0.0325 | | | | | |
| Thr3 | 0.3327 | 0.3728 | 0.2264 | 0.068 | | | | | | |
| Thr4 | 0.254 | 0.333 | 0.25 | 0.1305 | 0.0324 | | | | | |
| Tyr2 | 0.6723 | 0.0847 | 0.243 | | | | | | | |
| Val4 | 0.3398 | 0.2077 | 0.3102 | 0.0852 | 0.0571 | | | | | |
| Val5 | 0.2745 | 0.1968 | 0.2209 | 0.1905 | 0.0675 | 0.0499 | | | | |

*Table S8:* ***Mass Isotopomer Distribution (MID) results for the E. coli model with MetAMDB atom mappings (see Table S3).***

| Mass Isotopomer | M0 | M1 | M2 | M3 | M4 | M5 | M6 | M7 | M8 | M9 |
|---|---|---|---|---|---|---|---|---|---|---|
| AKG5 | 0.1781 | 0.2673 | 0.2766 | 0.1839 | 0.0728 | 0.0213 | | | | |
| AcCoA | 0.5497 | 0.2209 | 0.2294 | | | | | | | |
| Ala2 | 0.5617 | 0.2153 | 0.223 | | | | | | | |
| Ala3 | 0.5295 | 0.2285 | 0.0452 | 0.1967 | | | | | | |
| Asp2a | 0.4995 | 0.2993 | 0.2012 | | | | | | | |
| Asp2b | 0.4995 | 0.2993 | 0.2012 | | | | | | | |
| Asp3 | 0.3156 | 0.3617 | 0.2292 | 0.0935 | | | | | | |
| Asp4 | 0.2447 | 0.3257 | 0.2443 | 0.1349 | 0.0504 | | | | | |
| Cit6 | 0.1157 | 0.2308 | 0.2688 | 0.2156 | 0.117 | 0.0424 | 0.0097 | | | |
| E4P | 0.5423 | 0.1861 | 0.0469 | 0.0956 | 0.1291 | | | | | |
| Glu4 | 0.2276 | 0.3181 | 0.2802 | 0.1341 | 0.0401 | | | | | |
| Glu5 | 0.1794 | 0.267 | 0.2762 | 0.1834 | 0.0726 | 0.0213 | | | | |
| Gly1 | 0.726 | 0.274 | | | | | | | | |
| Gly2 | 0.6531 | 0.1417 | 0.2052 | | | | | | | |
| Ile5a | 0.1737 | 0.2686 | 0.2782 | 0.1849 | 0.0732 | 0.0214 | | | | |
| Ile5b | 0.1737 | 0.2686 | 0.2782 | 0.1849 | 0.0732 | 0.0214 | | | | |
| Leu5 | 0.1787 | 0.2669 | 0.2749 | 0.1847 | 0.0728 | 0.0221 | | | | |
| Mal4 | 0.2412 | 0.3244 | 0.2511 | 0.1349 | 0.0485 | | | | | |
| Met4a | 0.1916 | 0.3436 | 0.2812 | 0.1468 | 0.0368 | | | | | |
| Met4b | 0.1916 | 0.3436 | 0.2812 | 0.1468 | 0.0368 | | | | | |
| Met5 | 0.1486 | 0.2939 | 0.2763 | 0.1779 | 0.0836 | 0.0198 | | | | |
| Phe2 | 0.7178 | 0.068 | 0.2142 | | | | | | | |
| Phe8a | 0.1696 | 0.1841 | 0.2227 | 0.1498 | 0.1243 | 0.0737 | 0.051 | 0.0177 | 0.007 | |
| Phe8b | 0.1696 | 0.1841 | 0.2227 | 0.1498 | 0.1243 | 0.0737 | 0.051 | 0.0177 | 0.007 | |
| Phe9 | 0.1612 | 0.1832 | 0.1608 | 0.1706 | 0.1307 | 0.093 | 0.0529 | 0.03 | 0.0113 | 0.0063 |
| Pyr3 | 0.5169 | 0.2342 | 0.0467 | 0.2023 | | | | | | |
| Ser2a | 0.5055 | 0.3183 | 0.1762 | | | | | | | |
| Ser2b | 0.6885 | 0.1014 | 0.2101 | | | | | | | |
| Ser2c | 0.5055 | 0.3183 | 0.1762 | | | | | | | |
| Suc4 | 0.2376 | 0.323 | 0.2581 | 0.1348 | 0.0465 | | | | | |
| Thr3 | 0.3157 | 0.3616 | 0.2291 | 0.0935 | | | | | | |
| Thr4 | 0.2448 | 0.3257 | 0.2442 | 0.1349 | 0.0504 | | | | | |
| Tyr2 | 0.7179 | 0.068 | 0.2141 | | | | | | | |
| Val4 | 0.3024 | 0.2428 | 0.3009 | 0.1013 | 0.0526 | | | | | |
| Val5 | 0.2844 | 0.2429 | 0.1959 | 0.1752 | 0.0554 | 0.0464 | | | | |

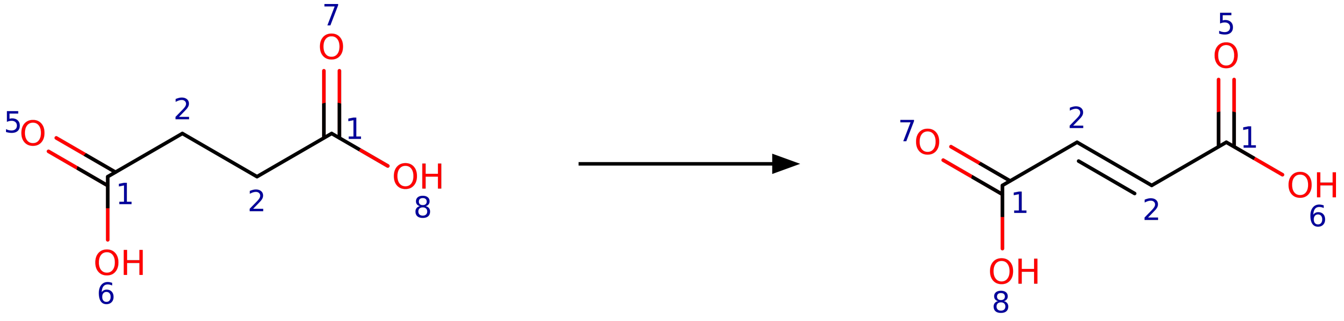


Succinate Fumarate

*Figure S1: Symmetry of the succinate dehydrogenase reaction*
